# Histamine originating from the BNST modulates corticostriatal synaptic transmission during early postnatal development

**DOI:** 10.1101/2023.10.19.563087

**Authors:** Ricardo Márquez-Gómez, Brenna Parke, Yasmin Cras, Sophie L. Gullino, Parry Hashemi, Tommas Ellender

## Abstract

The neuromodulator histamine regulates key processes in many regions of both the adult and developing brain including the striatum. However, striatal innervation by histaminergic afferents is very sparse making the physiological sources of histamine controversial. Here potential sources of striatal histamine were investigated during early postnatal development and specifically in the second postnatal week, in acute mouse brain slices. Firstly, a combination of whole-cell patch-clamp recordings and optogenetic stimulation demonstrates that during this period exogenously applied histamine modulates the intrinsic properties of developing D_1_ and D_2_ striatal spiny projection neurons (SPNs) as well as synaptic transmission at afferents coming from the mPFC and visual cortex. Secondly, immunohistochemistry for histamine reveals a brain region proximal and caudal to striatum densely innervated by histaminergic axons and corresponding to the oval nucleus of the bed nucleus of stria terminalis (ovBNST). Thirdly, direct electrical stimulation of the ovBNST leads to significant and detectable levels of histamine in the striatum, as assessed by both fast scan cyclic voltammetry and fluorescent histamine sensors. Lastly, electrical stimulation of the ovBNST nucleus, at frequencies mimicking active histaminergic neurons, can release sufficient levels of histamine to modulate excitatory synaptic transmission from mPFC onto striatal SPNs by acting at histamine H_3_ receptors. Together, these results provide evidence for the existence of the ovBNST as an extrastriatal source of histamine during early brain development and postulates a new view of the modus operandi of histamine in that it can cross anatomical boundaries and act as a paracrine neuromodulator.

**Significance statement:** Histamine is synthesized by neurons in the hypothalamic tuberomammillary nucleus (TMN) and released from their axons in many brain regions controlling key physiological processes. When dysregulated this can result in neurological and neurodevelopmental disorders such as Tourette’s syndrome and OCD. To understand the physiological roles for histamine and to facilitate the generation of new therapeutic interventions it is key to define the sources of histamine and its mode of action. Here we provide evidence, using the developing striatum as an exemplar, that sources of histamine can lie beyond anatomical boundaries with histamine acting as a paracrine neuromodulator. This also has potential implications for our mechanistic understanding of deep brain stimulation of the BNST in treating severe Tourette’s syndrome and OCD.

## Introduction

The neuromodulator histamine is synthesized by neurons located in the tuberomammillary nucleus (TMN) of the hypothalamus which send wide-spread axonal projections and release histamine throughout the central nervous system (<u>Haas</u> <u>& Panula, 2003</u>; <u>Haas et al., 2008</u>; <u>Lin et al., 2023</u>). Although classically assumed to mainly regulate the sleep-wake cycle, the histaminergic system is now thought to have many more diverse physiological roles in a multitude of brain regions and stages of life (<u>Schwartz et al., 1991</u>; <u>Takahashi et al., 2006</u>; <u>Panula et al., 2014</u>; <u>Han et al., 2020</u>; <u>Lucaci et</u> <u>al., 2023</u>), and has been shown to be dysfunctional in various neurological and neurodevelopmental disorders (<u>Ercan-Sencicek, 2010</u>; <u>Shan et al., 2015</u>; <u>Carthy & Ellender,</u> <u>2021</u>; <u>Xu et al., 2022</u>; <u>Ma et al., 2023</u>). For example, Tourette’s syndrome (TS) is an early-onset disorder that is characterized by involuntary motor and vocal tics often comorbid with obsessive-compulsive disorder (OCD) and depression (<u>Robertson et al., 2017</u>) and parallel findings in patients and mouse models suggest that a disruption of the histaminergic system and reduced levels of histamine can be part of the etiology (<u>Ercan-Sencicek, 2010</u>; <u>Fernandez et al., 2012</u>; <u>Karagiannidis et al., 2013</u>; <u>Baldan et al., 2014</u>). Symptoms of TS/OCD are thought to arise from dysregulation in the development and function of the striatum and its cortical inputs (<u>Nordstrom & Burton, 2002</u>; <u>McCairn et al.,</u> <u>2009</u>; <u>Worbe et al., 2012</u>; <u>Ahmari et al., 2013</u>; <u>Worbe et al., 2015</u>; <u>Corbit et al., 2019</u>) altering their interactions with other brain regions and associated structures (<u>O’Connell &</u> <u>Hofmann, 2012</u>; <u>Albin, 2018</u>) and can be exacerbated by stress and anxiety (<u>Robertson,</u> <u>2000</u>).

The early onset of these disorders has led to an increased interest in understanding the physiological and pathological roles for histamine during early prenatal and postnatal stages (<u>Panula et al., 2014</u>; <u>Rapanelli et al., 2017a</u>; <u>Han et al., 2020</u>; <u>Valle-Bautista</u> <u>et al., 2021</u>) as well as its potential as a target for treatment (<u>Rapanelli & Pittenger, 2016</u>; <u>Zhang et al., 2020</u>; <u>Carthy & Ellender, 2021</u>). It is puzzling that although histamine-expressing neurons can be readily detected in the hypothalamic tuberomammillary nucleus (TMN) from the first postnatal week onwards in rodents (<u>Reiner, 1988</u>; <u>Panula</u> <u>et al., 2014</u>) many brain regions, including the striatum, do not contain detectable histamine-containing axons until much later in life and even then are rather sparse (<u>Vanhala et al., 1994</u>; <u>Ellender et al., 2011</u>; <u>Han et al., 2020</u>; <u>Lin et al., 2023</u>). For example, in striatum these start to only be seen in the second postnatal week and even then are infrequent and in many striatal regions seemingly absent (<u>Han et al., 2020</u>). This developmental period is critical for the maturation of striatal spiny projection neurons (SPNs) and their synapses (<u>Kozorovitskiy et al., 2012</u>; <u>Kozorovitskiy et al., 2015</u>; <u>Peixoto</u> <u>et al., 2016</u>; <u>Lieberman et al., 2018</u>; <u>Krajeski et al., 2019</u>) and exogenously applied histamine has been shown to modulate both intrinsic and synaptic properties of developing SPNs (<u>Han et al., 2020</u>). This raises the intriguing question what the physiological sources of histamine might be and how these are able to interact with striatum.

Here a brain region is described located proximal and medial-caudal to the developing striatum which contains a high density of histaminergic fibers. Using whole-cell patch-clamp recordings, optogenetic stimulation, fast scan cyclic voltammetry (FSCV) and histamine-sensitive fluorescent reporters we demonstrate that this region provides an extra-striatal source of histamine capable of modulating developing striatal SPNs and their synapses and acting as a paracrine neuromodulator capable of crossing anatomical boundaries. Interestingly this region also corresponds to that successfully targeted in the treatment of severe TS and OCD.

## Results

### Histamine modulates the intrinsic electrical properties of developing D_1_ and D_2_ SPNs

The density of histamine-containing axons in the striatum is very sparse to non-detectable during the first postnatal weeks of life (<u>Vanhala et al., 1994</u>; <u>Han et al., 2020</u>). Even in adulthood this does not reach the axonal density exhibited by other neuromodulators, such as dopamine or serotonin (<u>Ellender et al., 2011</u>; <u>Awasthi et al.,</u> <u>2021</u>; <u>Lin et al., 2023</u>). The sparsity of innervation seems incongruous with the many observations demonstrating high expression of histamine receptors as well as histaminergic modulation of striatal neurons and synapses from the first postnatal weeks onwards (<u>Pillot et al., 2002</u>; <u>Ellender et al., 2011</u>; <u>Rapanelli et al., 2016</u>; <u>Rapanelli et</u> <u>al., 2017b</u>; <u>Han et al., 2020</u>). Indeed, during this period of development exogenously applied histamine has been shown to act on histamine receptors to modulate both the excitability of striatal spiny projection neurons (SPNs) and the activity-dependent plasticity at corticostriatal synapses (<u>Han et al., 2020</u>). To better understand how histamine impacts young developing striatal SPNs we first investigated whether histamine acts similarly on the two main SPN subtypes found in the striatum: the D_1_ and D_2_ SPNs. Whole-cell patch-clamp recordings were made in the dorsomedial striatum during the second postnatal week i.e., postnatal day (P)9-15 mice, from both types of SPN using biocytin-containing internal solution and *post hoc* staining for PENK (D_2_ SPNs) and DARPP-32 (striatal marker) to separate D_1_ SPNs from D_2_ SPNs (**Figure 1A, B**). Recordings were made in aCSF containing the GABA receptor antagonists SR95531 (200 nM) and CGP52432 (1 μM) and were made from a total of 56 SPN of which 21 could be classified as D_1_ and 21 could be classified as D_2_ SPNs. Superfusion of histamine (10 µM) led to a significant hyperpolarization of the membrane potential (aCSF: -67.93 ± 0.50 mV; Histamine: -70.36 ± 0.54 mV; p=0.0021, paired *t*-test, **Figure 1C**) and this hyperpolarization was seen for both D_1_ SPNs (-68.05 ± 0.98 mV to -70.45 ± 0.81 mV; p=0.048, paired *t*-test, **Figure 1C**) and D_2_ SPNs (-68.45 ± 0.97 mV to -71.26 ± 1.31 mV; p=0.010, paired *t*-test, n = both 21, **Figure 1C**). Concomitant with a general hyperpolarization of the membrane potential we observed a reduction in the action potential frequency elicited by increasing positive current injections (at +100 pA; aCSF: 17.10 ± 1.18 Hz vs Histamine: 13.18 ± 1.49 Hz and at +80 pA; 14.45 ± 1.01 Hz vs Histamine: 10.66 ± 1.38 Hz, paired *t*-test, p=0.012 and 0.019, n = 56, **Figure 1D**).

**Figure 1:**
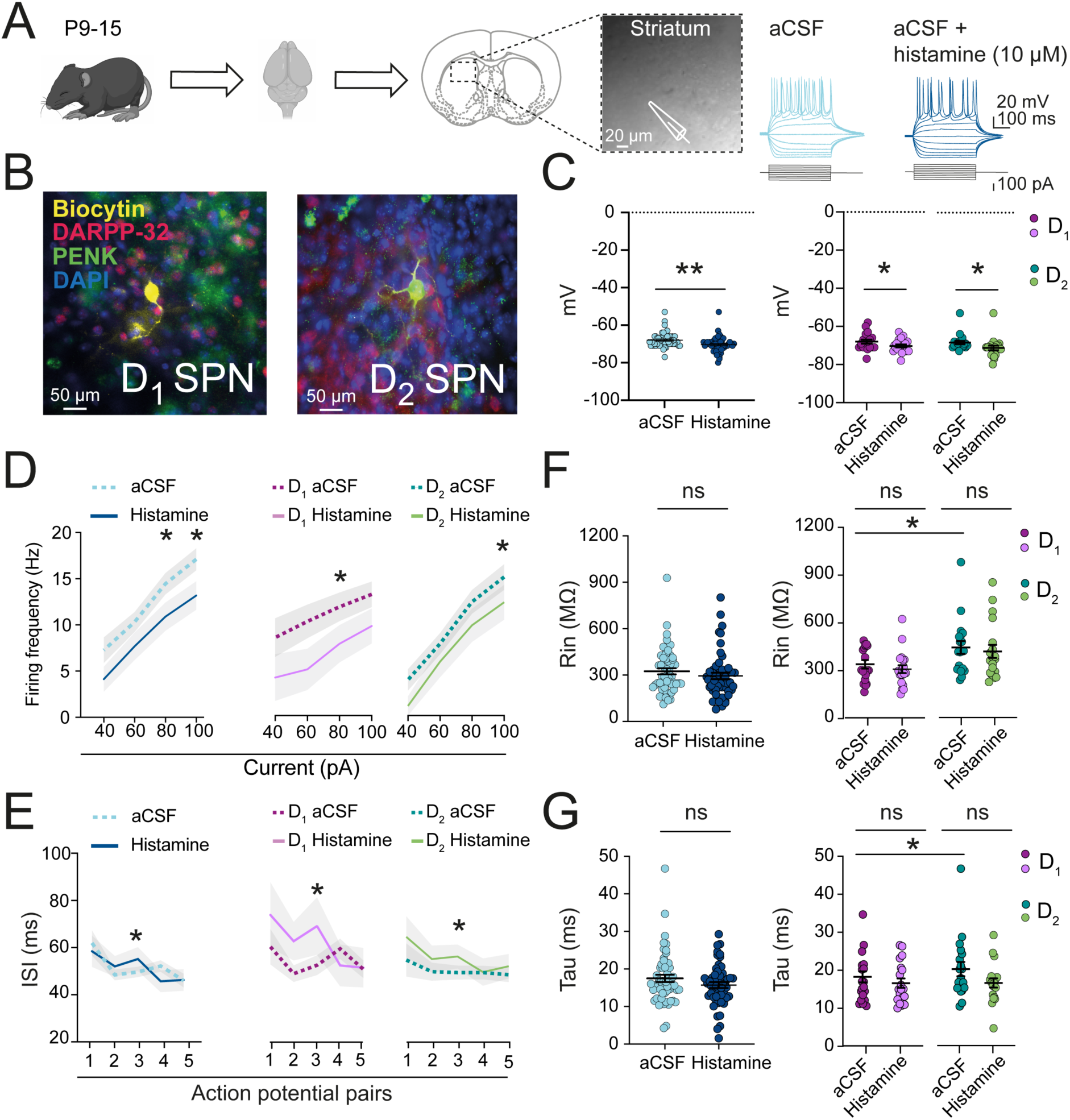
The electrical properties of developing D_1_ and D_2_ striatal SPNs are modulated similarly by histamine. (**A**) Postnatal day (P)9-15 C57Bl/6 mice were used to generate acute brain slices for whole-cell patch-clamp recordings of SPNs in the dorsomedial striatum to assess their intrinsic electrical properties in aCSF and during superfusion of aCSF containing histamine (10 µM). (**B**) Representative images of *post-hoc* immunostaining and classification of D_1_ and D_2_ SPNs. Co-localization of biocytin and PENK and DARPP-32 was indicative of D_2_ SPNs while biocytin and DARPP-32, but not PENK, co-localization was indicative of D_1_ SPNs. (**C**) Superfusion with aCSF containing histamine (10 µM) significantly hyperpolarized SPNs and this was seen for both D_1_ and D_2_ SPNs. (**D**) Histamine application led to a significant decrease in action potential frequency as seen in both D_1_ and D_2_ SPNs. (**E**) Histamine application shifted the interspike interval (ISI) to longer durations and was seen for both D_1_ and D_2_ SPNs. (**F**) The input resistance (Rin) of SPNs tends to decrease but was not significantly lowered by histamine. However, note the significantly higher Rin of the D_2_ SPNs under baseline conditions. (**G**) The membrane time constant or tau was not significantly changed by histamine but note the significantly longer membrane time constant of D_2_ SPN under baseline conditions. *p<0.05, **p<0.01

Similar reductions in action potential frequency were observed for both D_1_ SPNs (at +80 pA; aCSF: 14.17 ± 1.57 Hz vs Histamine: 9.31 ± 2.17 Hz, paired *t*-test, p=0.048, n = 21, **Figure 1D**) and D_2_ SPNs (at +100 pA; aCSF: 18.89 ± 1.41 Hz vs Histamine: 13.50 ± 2.02 Hz, paired *t*-test, p=0.029 n = 21, **Figure 1D**). The reduction in action potential frequency was consistent with observations of prolongations in the interspike interval (ISI) between pairs of action potentials upon histamine superfusion (pair 3; aCSF: 48.80 ± 1.68 ms vs Histamine: 56.35 ± 3.21 ms, paired *t*-test, p=0.035, n = 56, **Figure 1E**) and was seen for both D_1_ SPNs (pair 3; aCSF: 50.31 ± 2.79 ms vs Histamine: 72.90 ± 7.34 ms, paired *t*-test, p=0.039, n = 21, **Figure 1E**) and D_2_ SPNs (pair 3; aCSF: 48.94 ± 2.51 ms vs Histamine: 55.82 ± 3.88 ms, paired *t*-test, p=0.018, n = 21, **Figure 1E**). Lastly, although a general reduction in both the input resistance (Rin) (aCSF: 357.58 ± 18.97 MΩ and histamine: 330.90 ± 19.85 MΩ, p=0.25, paired *t*-test, n = 56, **Figure 1F**) and membrane time constant (Tau) (aCSF: 17.99 ± 0.89 ms vs Histamine: 16.23 ± 0.68 ms; p = 0.23, paired *t*-test, n = 56, **Figure 1G**) was seen, neither reached significance. However, it was observed that D_1_ SPNs had a consistently lower input resistance (D_1_: 320.88 ± 22.89 MΩ and D_2_: 428.56 ± 37.58 MΩ, p=0.023, independent-samples *t*-test, n = both 21, **Figure 1E**) and membrane time constant (D_1_: 17.60 ± 1.37 ms and D_2_: 20.32 ± 1.78 ms; p=0.042, paired *t*-test, n = both 21, **Figure 1G**) than D_2_ SPNs under baseline conditions consistent with previous observations and potentially resulting from differences in their maturational state (<u>Krajeski et al., 2019</u>). Overall, these results provide evidence of a significant modulatory role for histamine on developing striatal SPNs during the second postnatal week and suggest that both D_1_ and D_2_ SPNs are similarly affected during this period and these will therefore be grouped in subsequent experiments.

### Histamine modulates developing synaptic inputs onto SPNs coming from prefrontal and visual cortex

Histamine has been shown to modulate synaptic transmission at mature corticostriatal synapses as well as many other synaptic pathways (<u>Brown & Haas, 1999</u>; <u>Doreulee et</u> <u>al., 2001</u>; <u>Ellender et al., 2011</u>; <u>Zhang et al., 2020</u>) and more recently was shown to already modulate corticostriatal inputs during the first postnatal weeks when these are undergoing rapid maturation and are highly plasticity (<u>Peixoto et al., 2016</u>; <u>Valtcheva et</u> <u>al., 2017</u>; <u>Krajeski et al., 2019</u>; <u>Han et al., 2020</u>). Many of these studies used electrical and/or optogenetic stimulation of afferents coming from many different cortical regions simultaneously which might not all be similarly modulated. To study inputs coming from specific cortical regions here an optogenetic approach was employed based on local expression of the light-activatable channel ChR2, which also avoided potential problems with erroneous recruitment of non-cortical glutamatergic synapses (e.g. from thalamus (<u>Ellender et al., 2013</u>) or possible interactions with other modulatory afferents (e.g. dopamine) during electrical stimulation. This approach was used to isolate the input pathways from two distinct cortical regions, namely the visual cortex and medial prefrontal cortex (mPFC), which during the second postnatal week rapidly develop and especially for the latter is thought to be altered in Tourette’s syndrome and OCD (<u>George, 1992</u>; <u>Muller-Vahl et al., 2009</u>; <u>Ahmari, 2013</u>). To explore histaminergic modulation of these afferents AAV viral particles, containing the construct for ChR2 (hChR2(H134R)-eYFP) under the control of the CaMKIIa promotor (<u>Lee et al., 2010</u>), were injected into either the mPFC or visual cortex of P0-3 mice to transfect their constituent excitatory neurons. After allowing for sufficient time for ChR2-eYFP expression, acute striatal brain slices were made at P9-15 for whole-cell current-clamp recordings of striatal SPNs (**Figure 2A, B**). Robust expression of ChR2-eYFP was seen in medial striatum (<u>van Heusden et al., 2021</u>) and brief flashes of blue light (3 ms, 473 nm) elicited robust excitatory postsynaptic potentials (EPSPs) in recorded SPNs (**Figure 2B, C**). Optical activation of mPFC inputs generated EPSPs with a mean amplitude of 2.28 ± 0.03 mV (**Figure 2C** and **Table 1**) with superfusion of histamine (10 μM) causing a significant decrease in their amplitude (to 83.47 ± 12.80% of baseline, p=0.031; Wilcoxon test, n = 23, **Figure 2C, D**) duration and decay time (decay time to 84.65 ± 7.14% of baseline; p=0.022, Wilcoxon test, n=23, **Figure 2C, D** and **Table 1**). To investigate whether this histaminergic modulation of developing corticostriatal synapses is unique or biased towards afferents coming from mPFC the experiments were repeated for inputs coming from the visual cortex (**Figure 2A, B**). The visual cortex sends functional projections to striatum (<u>Khibnik et al., 2014</u>; <u>van</u> <u>Heusden et al., 2021</u>) and develops rapidly after eye-opening during the latter part of the second postnatal week in mice (<u>Shen & Colonnese, 2016</u>). Indeed, optical activation of visual cortex inputs also generated robust EPSPs with a mean amplitude of 4.28 ± 1.28 mV (**Figure 2C** and **Table 1**). Similarly to the observations for mPFC, histamine also decreased the amplitude of synaptic inputs coming from visual cortex (to 69.41 ± 6.85%, p=0.0019, Wilcoxon test, n=18, **Figure 2E**), as well as reducing their duration and decay time (decay time to 75.72 ± 11.25%, p=0.01, Wilcoxon test, n=18, **Figure 2C** and **Table 1**). These results demonstrate that during the second postnatal week, histamine can negatively modulate the striatal excitatory inputs onto SPNs coming from both mPFC and visual cortex. This, together with the previous findings on the histaminergic modulation of intrinsic properties, supports the idea that histamine is an active neuromodulator during this important developmental period and is able to control diverse processes at striatal SPNs.

**Figure 2.**
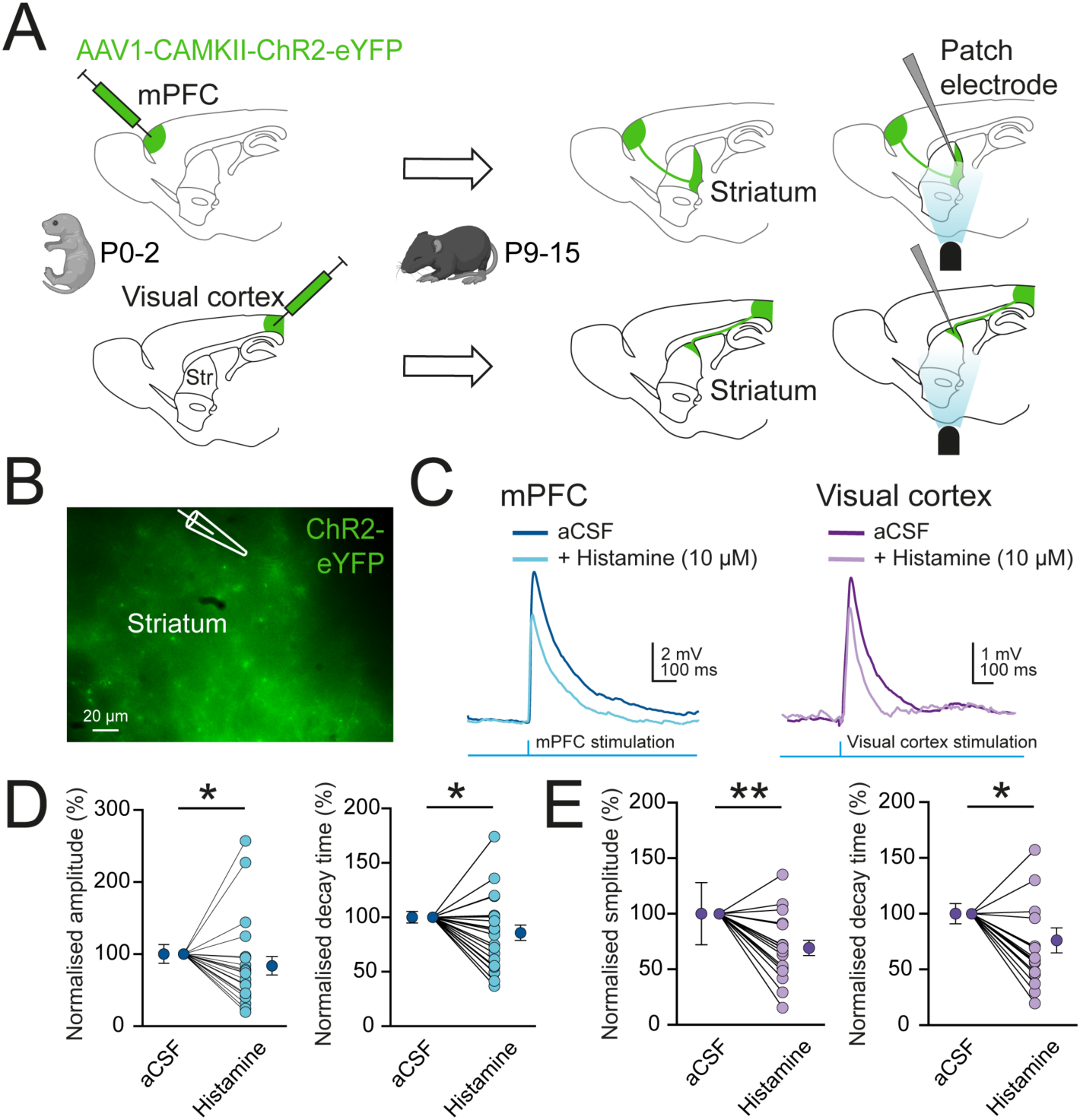
Developing corticostriatal synapses from mPFC and visual cortex onto SPNs are modulated by histamine. (**A**) Experimental diagram; P0-2 pups were injected with AAV1-CAMKII-ChR2-eYFP in the medial prefrontal cortex (mPFC, top) or visual cortex (bottom) and after allowing for sufficient time for ChR2-eYFP expression whole-cell current-clamp recordings were made from SPNs in acute striatal slices during the second postnatal week. (**B**) Example image of striatum from a mPFC injected mouse demonstrating dense ChR2-eYFP-expressing axons in dorsomedial striatum. (**C**) Brief pulses of blue light (3 ms) were used to stimulate afferents from mPFC and visual cortex and elicited robust excitatory postsynaptic potentials (EPSPs) as recorded from SPNs. Representative traces of EPSPs obtained from optogenetic stimulation during baseline conditions and in the presence of histamine. (**D**) Both the amplitude and decay time of the mPFC-evoked EPSPs were negatively modulated by histamine (10 µM) which was also seen for (**E**) visual cortex-evoked EPSPs. *p<0.05, **p<0.01

**Table 1:** EPSP properties of mPFC and visual cortex inputs onto striatal SPNs.

|  | aCSF | Histamine | p-value |
| --- | --- | --- | --- |
| **Medial prefrontal cortex (mPFC)** |  |  |  |
| Amplitude (mV) | 2.63 ± 0.32 | 2.09 ± 0.40 | 0.031 |
| Duration (ms) | 262.3 ± 11.73 | 229.7 ± 14.69 | 0.035 |
| Rise Time (ms) | 9.14 ± 0.47 | 9.30 ± 1.01 | 0.99 |
| Decay Time (ms) | 117.30 ± 6.14 | 96.51 ± 7.64 | 0.022 |
| Time to peak (ms) | 18.28 ± 0.94 | 18.62 ± 2.03 | 0.99 |
| **Visual cortex** |  |  |  |
| Amplitude (mV) | 4.28 ± 1.28 | 2.98 ± 0.81 | 0.0019 |
| Duration (ms) | 234.70 ± 21.47 | 163.10 ± 17.17 | 0.02 |
| Rise Time (ms) | 10.71 ± 1.14 | 12.14 ± 2.67 | 0.67 |
| Decay Time (ms) | 107.00 ± 10.20 | 69.40 ± 7.35 | 0.01 |
| Time to peak (ms) | 21.42 ± 2.29 | 24.28 ± 5.34 | 0.67 |

Data are given as mean ± SEM, statistical comparisons by paired Wilcoxon *t*-test

### Histamine-containing axons form a dense axonal plexus at the caudal border of striatum

The fact that exogenously applied histamine modulates young striatal neurons and synapses raises the question what the physiological sources for striatal histamine might be. Puzzlingly, numerous observations would suggest that histamine-containing axons are sparse to undetectable in many brain regions, including the striatum, during both early development as well as in adulthood (<u>Ellender et al., 2011</u>; <u>Yu et al., 2015</u>; <u>Han</u> <u>et al., 2020</u>; <u>Lin et al., 2023</u>). To explore in detail the rostro-caudal distribution of histamine-containing axons in the developing brain fixed sagittal brain sections from P9-10 mice were immunolabeled for histamine (**Figure 3A - C**). Firstly, as expected a dense region of histamine-positive neurons and fibers was observed in the hypothalamic area corresponding to the anatomical location of the tuberomammillary nucleus (TMN, **Figure 3A**) (<u>Haas & Panula, 2003</u>) and little to no histamine-containing axons were observed in striatum (<u>Han et al., 2020</u>).

**Figure 3:**
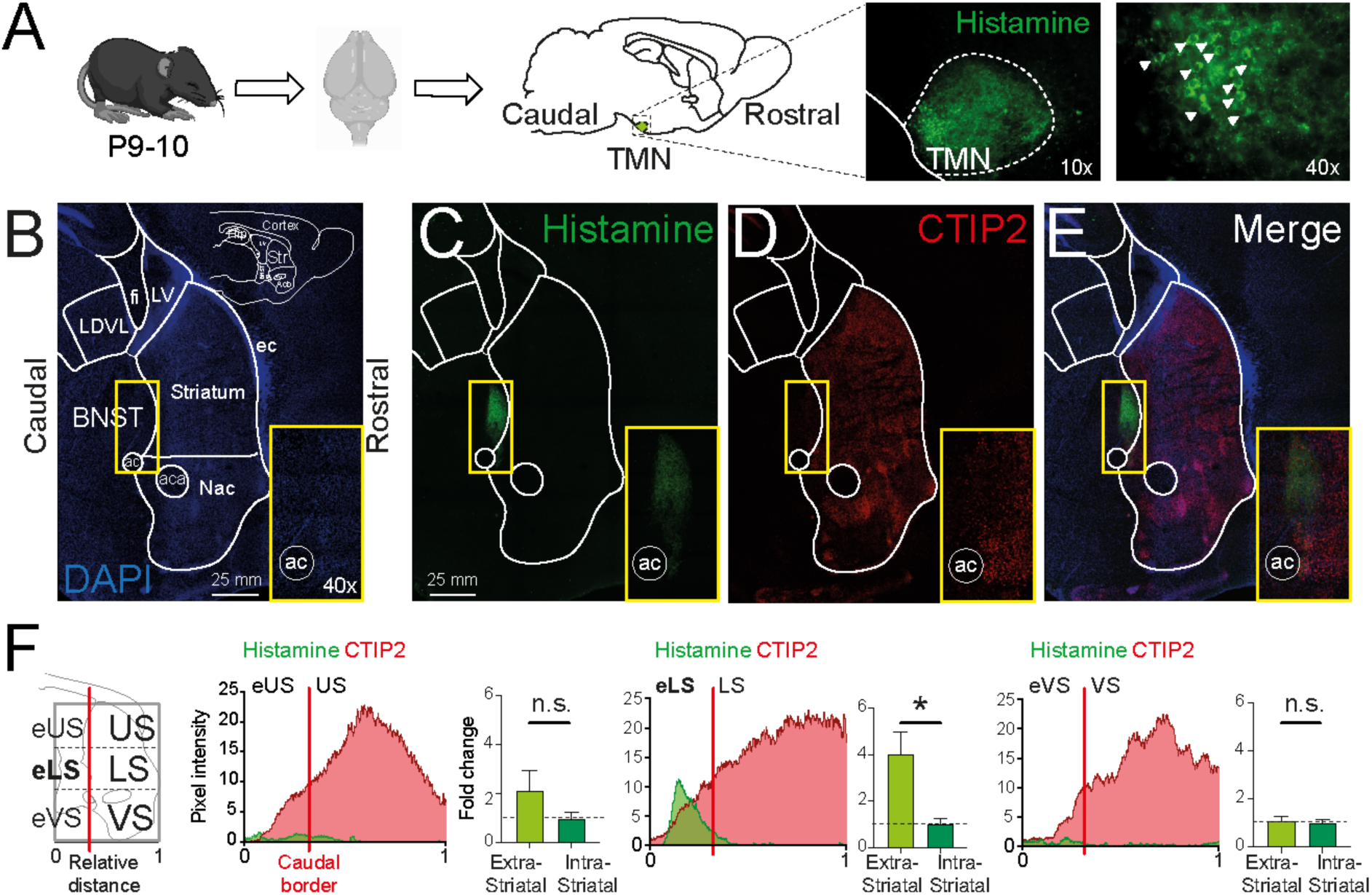
Histamine fibers form a dense axonal cluster located caudal from striatum. (**A**) Experimental approach; sagittal brain sections of P9-10 C57Bl/6 mice were labelled for histamine. Densely packed histaminergic neurons (white arrow heads) and fibers were found in the hypothalamic tuberomammillary nucleus (TMN). (**B**) Sagittal brain section labelled with the nuclear dye DAPI with brain regions indicated that are observable at this plane of cutting (lateral: 0.96 mm, Franklin, 2007). Note the orientation of sections with rostral to the right and caudal to the left. (**C**) Histamine staining showed a dense axonal plexus at the caudal border of striatum (yellow rectangle: higher magnification). (**D**) Labelling for CTIP2 reveals this region is somewhat positive but with much lower levels of expression as compared to striatum. (**E**) Merge of histamine, CTIP2 and DAPI staining. (**F**) To dissect histamine fiber distribution further, the striatum and more caudal regions were divided in 6 distinct areas comprising striatal (US, LS, VS) an extra-striatal (eUS, eLS and eVS) areas. The red line represents the anatomical border of the caudal striatum. Analysis of the intensity of histamine labelling across the striatal and extra-striatal regions revealed the strongest signal (both in the pixel intensity plots derived from one representative brain section, as well as the fold change plots representing histamine signal normalized to intrastriatal histamine averaged for all data) mainly located to the external portions of the striatum and specifically the eLS (middle plots) with significantly lower levels of expression seen in all other regions. LV=lateral ventricle, fi=fimbria, LDVL=laterodorsal nucleus of the thalamus, BNST=bed nucleus of stria terminalis, NAc=nucleus accumbens, ac=anterior commissure, aca=anterior arm of anterior commissure, ec=external capsule. * p<0.05

However, when exploring the borders of striatum an unexpected dense plexus of histamine-containing axons was observed close to the caudal border of striatum located ventrally from the lateral ventricle (LV) and dorsally from the anterior commissure (*ac*; **Figure 3C**). This cluster of histamine-positive axons co-localized with a region expressing low levels of the striatal marker CTIP2 (**Figure 3D, E**) (<u>Arlotta et</u> <u>al., 2008</u>). Although CTIP2 is an often used striatal marker, it is also expressed in other brain regions and neurons, especially those that originate from the embryonic lateral ganglionic eminence (LGE) (<u>Arlotta et al., 2008</u>; <u>Tinterri et al., 2018</u>). Next the precise anatomical location and boundaries were delineated using a detailed investigation across multiple sagittal brain sections, using the edge of the *ac* as a reliable marker indicating the caudal limit of the striatum and the external capsule (*ec*) as the rostral start of striatum (**Figure 3B)**. Using these landmarks, and assisted with DAPI staining and the Paxinos mouse brain atlas (<u>Franklin, 2007</u>), the striatum was segmented into three regions: an Upper (US), Lower (LS) and Ventral (VS) Striatum (**Figure 3B, F**). In addition, as the *ac* was taken as the border of the caudal striatum, more caudal regions were referred to as external US (eUS), external LS (eLS) and external VS (eVS) and all corresponding to extra-striatal regions. Following this segmentation, most histamine-expressing fibers were observed in a CTIP2-positive region which exhibited a distinct expression profile from that of the striatum. Indeed, note the reduced CTIP2 signal corresponding to regions when the histaminergic signal is high (**Figure 3C, D** and **F**), suggesting the region where the histamine fibers are located is distinct and lies beyond the anatomical caudal limit of the striatum. Quantification of these images reveals that the enriched histamine axonal region was more likely to be found in the eLS (67.21 ± 7.41%) as compared with the eUS or eVS (18.77 ± 2.34% and 12.66 ± 1.11%) and detection of histamine fibers within any sector of the striatum was on average low and limited to regions very close to the caudal striatum in agreement with previous findings (<u>Han et al., 2020</u>). Careful study of the eLS region in our brain sections in conjunction with the Paxinos brain atlas (<u>Franklin, 2007</u>) would suggest this anatomical region corresponds to the bed nucleus of stria terminalis or BNST.

### The BNST oval nucleus contains dense histaminergic innervation

Anatomical analysis suggested that the histamine fibers were located in a region at the caudal border of striatum and likely corresponded to the BNST (<u>Franklin, 2007</u>). The BNST forms part of the brain’s limbic system and consists of a complex collection of nuclei thought to regulate a variety of stress and anxiety related behaviors (<u>Lebow &</u> <u>Chen, 2016</u>). These subnuclei include the oval nucleus of the BNST (ovBNST) whose activity is thought to promote anxiety-related behaviors with more ventro-caudal regions such as the ventral matrix of the BNST (BNSTvm) having more anxiolytic roles (<u>Dong et al., 2001</u>; <u>Kim et al., 2013</u>).

As the histaminergic axonal plexus appeared highly localized, we next explored whether it was constrained to a particular nucleus of the BNST. Previous work has shown that ovBNST neurons express high levels of the marker PKC-8 (<u>Tinterri et al.,</u> <u>2018</u>; <u>Wang et al., 2019</u>). Labelling for PKC-8 in our sections indeed revealed a small distinct region located at the caudal border of the striatum (**Figure 4A**). Interestingly, the histaminergic axonal plexus, highly co-localized with this PKC-8 positive region (**Figure 4A, B**). Quantification over multiple sections revealed that 90.79 ± 2.15% of histamine fibers co-localized with the PKC-8 positive region corresponding to the ovBNST, with the remainder of positive fibers (9.21 ± 2.15%) found outside of this region (p=0.0028, paired *t*-test, n = 3 mice, **Figure 4C**). These results suggest the presence at an early developmental age of a dense histamine axonal plexus constrained to the ovBNST nucleus.

**Figure 4.**
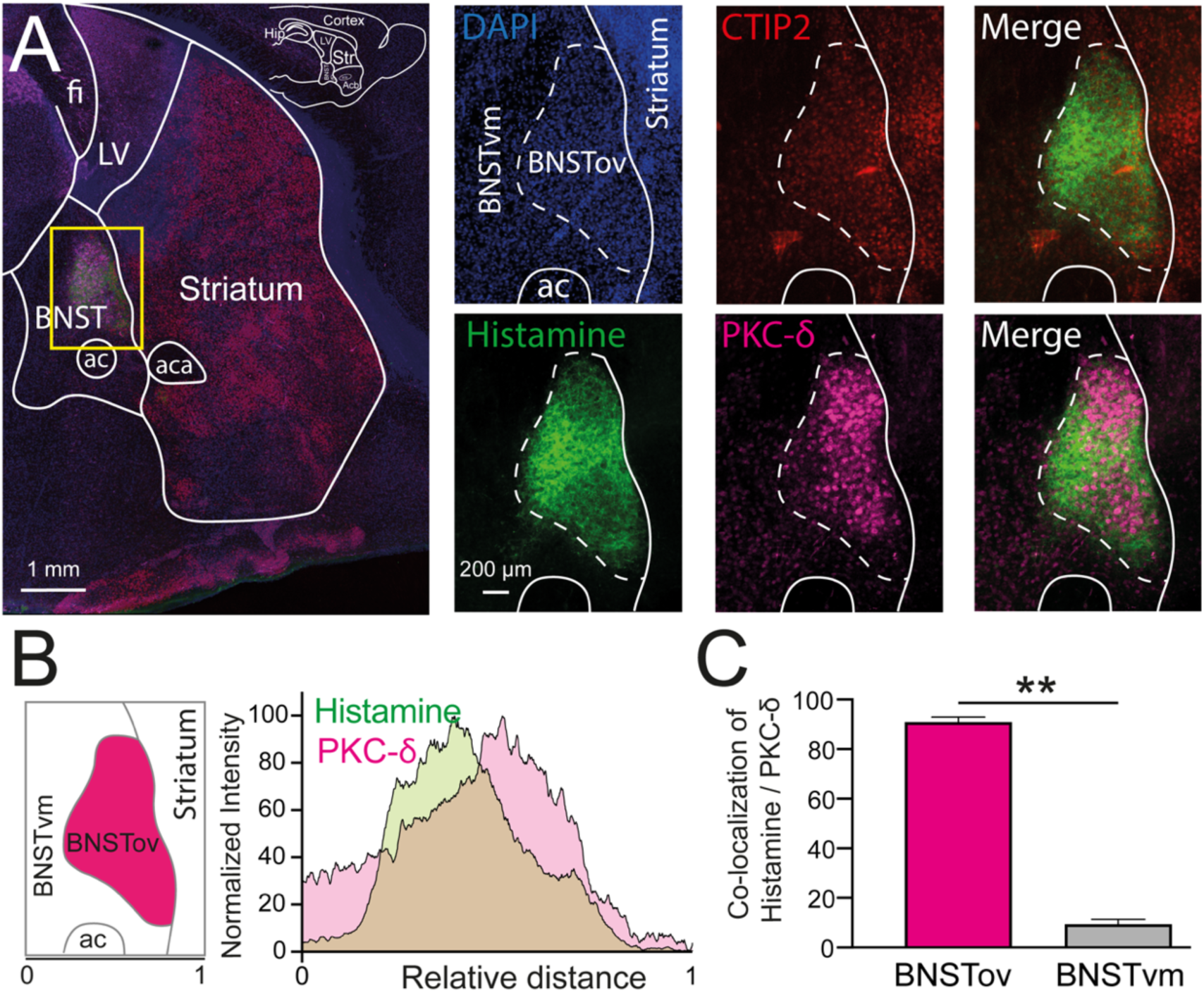
A dense histamine axonal plexus is found in the oval nucleus of the bed nucleus of stria terminalis (ovBNST) (**A**) Representative image of a sagittal brain section labelled for histamine revealing the location of the histamine positive axons in the BNST. Sections were also co-labelled for the striatal marker CTIP2 and the ovBNST marker PKC-8. Note the overlap between PKC-8 and histamine staining. (**B**) Quantification of the intensity of histamine and PKC-8 labelling in the brain section and region indicated in the yellow rectangle in A. (**C**) Quantification across multiple brain sections and mice showed that over 90% of the histamine staining is constrained to the PKC-δ positive region corresponding to ovBNST. LV=lateral ventricle, fi=fimbria, BNST=bed nucleus of stria terminalis, ac=anterior commissure, aca=anterior arm of anterior commissure, BNSTvm=ventromedial nucleus of the bed nucleus of stria terminalis. ** p<0.01

### Significant levels of histamine can be released from ovBNST and is detectable within striatum

Having demonstrated that the ovBNST contains a dense histaminergic axonal plexus we next explored whether this could represent a possible physiological source of striatal histamine. Histamine concentrations in the brain have classically been investigated using techniques such as microdialysis and high-performance liquid chromatography which have suggested that extracellular histamine can be detected for long periods of time after electrically evoked release in adult rodents and is detectable for minutes (<u>Cumming, 1991a</u>). More recent techniques based on fluorescent sensors also support this extended life-time of extracellular histamine (<u>Dong et al.,</u> <u>2023</u>). This contrasts with many other monoamines including dopamine, which has a half-life ranging in the hundreds of milliseconds and highly spatially restricted actions (<u>Rice & Cragg, 2008</u>; <u>Salinas et al., 2016</u>; <u>Sulzer et al., 2016</u>; <u>Hamid et al., 2021</u>). Giving the high density of histamine containing axons close to the caudal border of the striatum and the demonstrated extended lifetime once released, we next investigated whether the ovBNST could act as an extra-striatal source of histamine.

Two approaches were used to investigate whether this might be the case. Firstly, fast cyclic voltammetry measurements were made with carbon fibers placed in the striatum close to the ovBNST. The ovBNST was briefly stimulated (20 Hz for 1 second) using Tungsten electrodes and possible histaminergic transients detected from carbon fibers (distance between stimulating and recording electrodes: 497 ± 51 μm, n=6 brain sections from 3 mice). These experiments demonstrated that ovBNST stimulation can elicit both a fast dopaminergic transient and a slightly delayed and slower histaminergic transient as detected in the striatum (**Figure 5A**). To validate and verify these cyclic voltammograms obtained from brain sections the carbon fiber electrodes were also placed in stirred beakers of oxygenated recording aCSF containing low concentrations of histamine (100 nM) and dopamine (100 nM; **Figure 5B**) which confirmed that histamine and dopamine were detected. On average we find that electrical stimulation led to a significant increase of both dopamine (45.3 ± 5.6 nM, paired *t*-test, p=0.0002) and histamine (178.5 ± 57.7 nM, paired *t*-test, p=0.009) which peaked at respectively 5.6 ± 0.07 s and 7.0 ± 0.2 s after stimulation (p=0.0016, paired *t*-test) with histamine lingering for significantly longer in the tissue (dopamine: 6.6 ± 0.4 s and histamine: 18.7 ± 3.7 s, p=0.019, paired *t*-test, n=6 brain sections from 3 mice, **Figure 5C**). Secondly, to further confirm that stimulation of ovBNST can generate a significant histamine transient in the striatum recently developed GRAB_HA_ fluorescent sensors were used (<u>Dong et al., 2023</u>). Here plasmid DNA containing the HA1h variant of the GRAB_HA_ sensor was injected into the ventricles of developing mouse embryos and delivered to neuronal progenitors in the embryonic LGE using *in utero* electroporation (<u>van Heusden et al., 2021</u>). This allowed for the labelling of postnatal striatal SPNs with the HA1h GRAB_HA_ sensor which is endogenously fluorescent (<u>Dong et al., 2023</u>) (**Figure 5E**). Acute coronal striatal sections were made from electroporated pups at P9-15 of age and sections were selected that contained the ovBNST as well as SPNs located near the ovBNST expressing the HA1h GRAB_HA_ sensor (**Figure 5E, F**). To first confirm that the sensor is responsive to histamine these HA1h GRAB_HA_ sensor-expressing SPNs were imaged for brief periods of time (1-2 seconds) while being illuminated with a wide-field 473 nm LED both before and during superfusion with histamine (10 µM). This revealed that indeed this method of expression in striatal neurons led to functional GRAB_HA_ sensors which exhibited a significant increase in fluorescence in the presence of histamine (0.30 ± 0.09, *ΔF*/*F_0_*, p=0.049, one sample *t*-test, n = 4 cells, 3 mice) comparable to that seen using other methods of delivery of the sensor (<u>Dong et al., 2023</u>) (**Figure 5G**).

**Figure 5.**
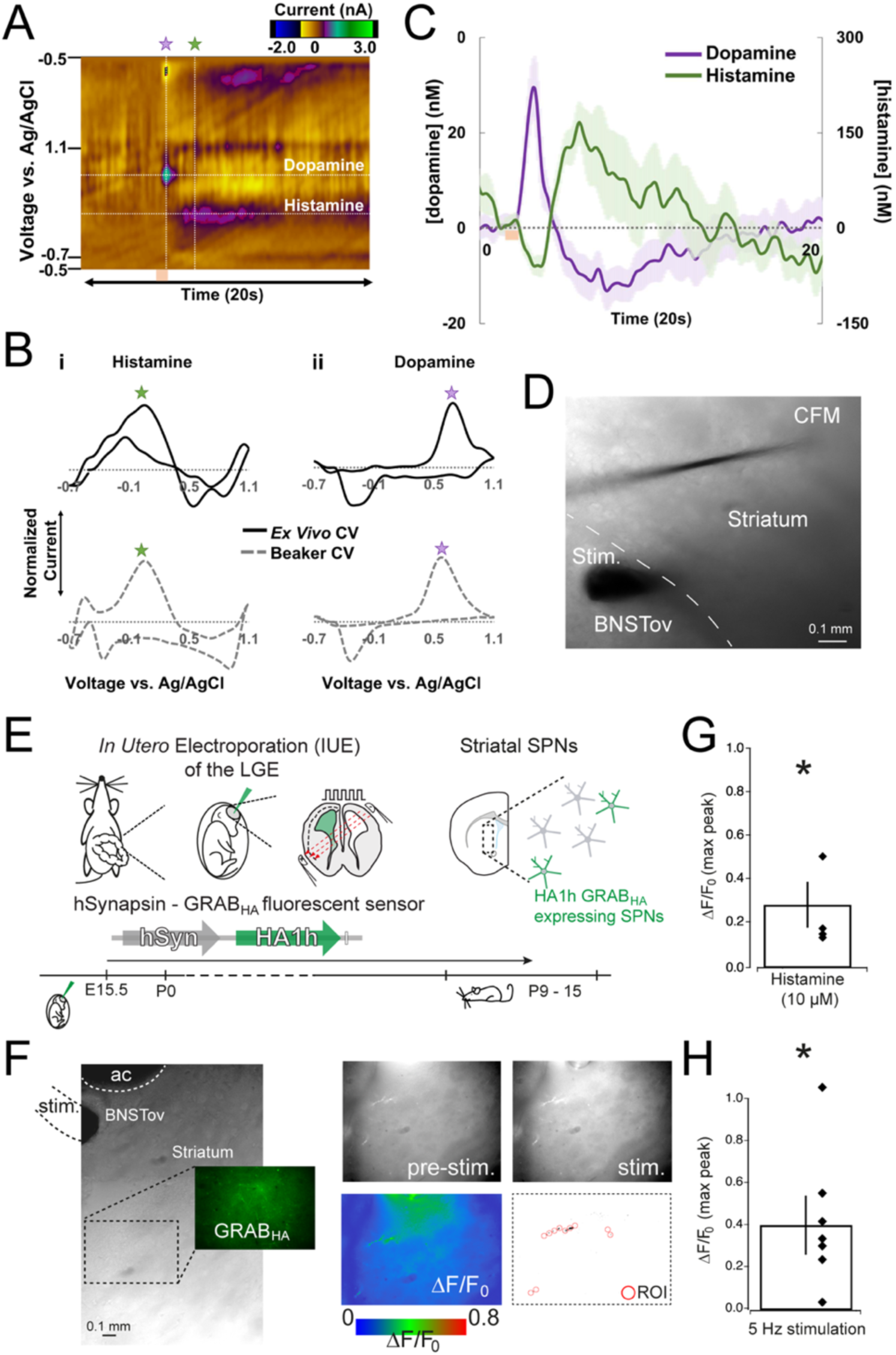
Electrical stimulation of the ovBNST leads to significant and detectable levels of histamine in striatum. (**A**) FSCV recordings in the striatum during ovBNST stimulation. Representative color plot of a stimulated response signal at the CFM where the first vertical line (purple star) reflects dopamine release, and the second vertical line (green star) reflects histamine release. (**B**) The cyclic voltammograms (CVs) from the brain (**Bi** and **Bii**, top) were verified by introducing low concentrations of histamine (**Bi**, bottom) and dopamine (**Bii**, bottom) in a stirred beaker of recording aCSF (30 ⁰C) bubbled with carbogen. (**C**) Bandpass filtered (2^nd^ order Butterworth with 0.01 to 1.5 Hz cutoff frequencies) current vs. time (IT) traces for dopamine and histamine, averaged (n=6 stimulations from 3 mice) with error reported as standard error of the mean. (**D**) An illustrative representation of the experimental setup where the CFM was placed in the striatum close to the stimulation electrode placed in the ovBNST. (**E**) *In utero* electroporation was used to deliver the histamine sensitive GRAB_HA_ sensor to embryonic progenitors of the LGE resulting in expressing of the sensor in postnatal striatal SPNs. Acute coronal striatal slices were made and checked for endogenous fluorescence of the GRAB_HA_ sensor in the somata and dendrites of transfected SPNs. (**F**) Experimental setup: a bipolar stimulation electrode was placed in the ovBNST, and histamine release was measured from changes in fluorescence of the GRAB_HA_ sensor expressed by SPNs located close to the ovBNST. Note the increase in fluorescence during stimulation as a change in *ΔF/F_0_* (**G**) Superfusion of slices with histamine (10 µM) led to a significant increase in the *ΔF/F_0_* signal. (**H**) Electrical stimulation of the ovBNST produced a significant increase in *ΔF/F_0_* as measured in striatum from GRAB_HA_ sensor expressing neurites within chosen regions of interest (ROI), suggesting that histamine released from the ovBNST migrates into the striatum.

Following a similar approach as in the FSCV experiments, we next placed a stimulating electrode in the ovBNST and induced histamine release by applying 30 seconds of electrical stimulation at a frequency of 5 Hz reflecting naturalistic activity patterns of TMN histaminergic neurons (<u>Biggs & Johnson, 1980</u>; <u>Haas & Panula, 2003</u>). Changes in the fluorescent signal were detected in neurites from GRAB_HA_ sensor expressing SPNs located close to the ovBNST (distance between stimulating electrode and the center of expressing neurites: 512 ± 52 μm, n=7 brain sections from 6 mice). This revealed a robust increase in the fluorescence (0.56 ± 0.18 *ΔF*/*F_0_*, p=0.028, one sample *t*-test, n = 7 cells, 6 mice). These two experiments demonstrate that histamine released from the ovBNST can cross anatomical barriers and migrate into the striatum, acting as an extra-striatal source of histamine.

### Histamine released from ovBNST modulates transmission at mPFC corticostriatal axons

These results demonstrate that by the second postnatal week the intrinsic electrical properties of D_1_ and D_2_ SPNs and their cortical inputs from mPFC and visual cortex can be actively modulated by histamine, that a dense histamine-positive axonal plexus can be found close to striatum corresponding to the ovBNST, and that electrical stimulation of the ovBNST can result in detectable histamine transients within the striatum as measured with CFM and GRAB_HA_ sensors. We next investigated whether these electrically evoked histamine transients were sufficient and large enough to modulate striatal physiology and specifically corticostriatal synaptic transmission. These experiments combined simultaneous optogenetic activation of mPFC afferents, electrical stimulation of ovBNST and whole-cell currents-clamp recordings of SPNs in medial striatum (**Figure 6A**). The striatum also receives direct inputs from the BNST (<u>Smith et al., 2016</u>), and to prevent direct axonal stimulation of these projections, a thin and precise cut was made between the BNST and the striatum (see methods, **Figure 6B**). Optogenetic stimulation of mPFC axons with brief light pulses (3 ms, 473 nm) led to reliable and stable EPSPs as detected from patched SPNs (**Figure 6C**). After a 5-minute baseline period the ovBNST was electrically stimulated at 5 Hz frequency for 3 minutes reflecting naturalistic activity pattern of TMN histaminergic neurons (<u>Biggs &</u> <u>Johnson, 1980</u>; <u>Haas & Panula, 2003</u>). Following electrical stimulation of the ovBNST the optogenetic stimulation was resumed for a further 10 minutes. Analysis of the normalized EPSP amplitude revealed a significant reduction in their amplitude seen consistently around 4-5 minutes after electrical stimulation (aCSF baseline: 100.40 ± 2.62%, aCSF post stim: 47.91 ± 3.23%; one-Way ANOVA p=<0.0001, n = 10 neurons/ 5 mice, **Figure 6C** and **6D**). During the entire recording both the membrane potential (p=0.12; Wilcoxon test, 5 min) and input resistance (p=0.06, Wilcoxon test, 5 min) did not significantly change (**Figure 6C**). As the BNST is also innervated by many other neuromodulatory systems, including dopamine and serotonin (<u>Lebow & Chen, 2016</u>; <u>Chiang et al., 2020</u>; <u>Donner et al., 2020</u>), and these have the potential to contribute to these observations the experiment was repeated in the presence of the H_3_R antagonist thioperamide (10 µM). Under these conditions, the reduction in optogenetically evoked mPFC-striatal EPSP amplitude was significantly reversed (aCSF + thioperamide: to 89.60 ± 3.09%; one-Way ANOVA, p=0.98, n = 10 neurons / 5 mice, **Figure 6C** and **6D**), suggesting that for the most part our results can be explained by histamine acting at H_3_R. Together, these results support the idea that the ovBNST can act as an extra-striatal source of histamine and that released histamine can act in a paracrine manner and can modulate mPFC-striatal inputs during the early developmental periods studied.

**Figure 6.**
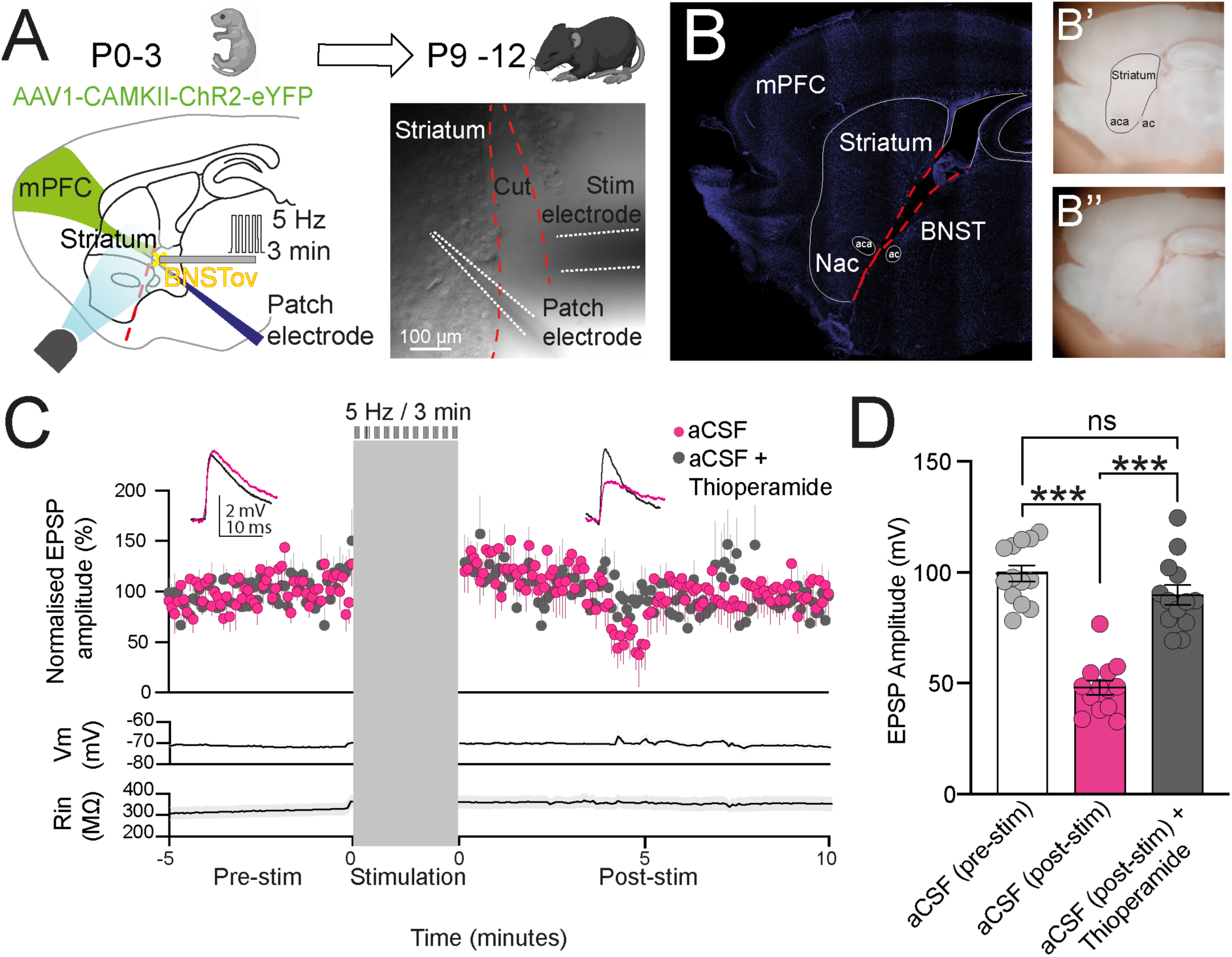
Histamine released from ovBNST can modulate corticostriatal synaptic transmission. (**A**) Experimental design: P3 C57Bl/6 mice were injected in the mPFC with AAV1-CAMKII-ChR2-eYFP and sagittal brain slices were generated from mice between P9-12. An electrical stimulation electrode was placed in the ovBNST and whole-cell patch-clamp recordings were made from close-by striatal SPNs. A thin cut was made between the ovBNST and striatum to sever direct synaptic connections between ovBNST and striatum. (**B**) Image of a fixed sagittal brain section stained with DAPI illustrating the location of the cut (left) as well as *in situ* in fresh tissue (right) before B’ and after B’’ the cut. (**C)** Electrical stimulation of ovBNST (3 min @ 5Hz) caused a transient decrease in the amplitude of the evoked EPSPs starting around 4-5 minutes after stimulation (pink circles). Superfusion of aCSF containing the H_3_R antagonist thioperamide (10 µM) significantly attenuated this reduction in EPSP amplitude (grey circles). The input resistance (Rin) and membrane voltage (Vm) were monitored during the recording and did not significantly alter. (**D**) Quantification of the EPSP amplitude observed before and after BNST stimulation (pink) and the attenuation of histaminergic modulation by blocking the H_3_R with thioperamide (grey). mPFC=medial prefrontal cortex, BNST=bed nucleus of stria terminalis, NAc=nucleus accumbens, ac=anterior commissure, aca=anterior arm of the anterior commissure. *** p<0.001

## Discussion

Our results suggest that the oval nucleus of the bed nucleus of stria terminalis (ovBNST) can provide an extrastriatal source of histamine during early postnatal development when histaminergic innervation of striatum itself is sparse to undetectable. We first demonstrate that striatal neurons during this developmental period are responsive to histamine by showing that both the intrinsic properties of D_1_ and D_2_ striatal spiny projection neurons and their cortical inputs coming from mPFC and visual cortex are actively modulated by exogenously applied histamine. Next using immunohistochemistry for histamine, various further molecular markers, and detailed anatomical analysis we find a region proximal to striatum that is densely innervated by histaminergic afferents and corresponds to the ovBNST. We then go on to show that direct electrical stimulation of the ovBNST leads to significant and relatively prolonged histamine transients in striatum as detected using both carbon fiber voltammetry as well as genetically expressed fluorescent histamine sensors. Lastly, by combining electrical stimulation of the ovBNST and optogenetic stimulation of mPFC afferents we are able to show that these histamine transients are sufficient and large enough to transiently modulate corticostriatal synaptic transmission by acting at histamine H_3_ receptors. Together, these results suggest a novel view on the mode of action of this important neuromodulator in that histamine is able to cross anatomical boundaries and act as a paracrine neuromodulator. This is demonstrated through direct electrical stimulation of the ovBNST which interestingly is also an efficacious target for deep brain stimulation in disorders related to dysregulated histamine.

It is puzzling that striatal innervation by histaminergic afferents is sparse or undetectable in the first postnatal weeks (<u>Vanhala et al., 1994</u>; <u>Han et al., 2020</u>) and even in adulthood is not very dense, as compared to other neuromodulators such as dopamine (<u>Ellender et al., 2011</u>; <u>Awasthi et al., 2021</u>; <u>Lin et al., 2023</u>). Close examination of sagittal sections stained for histamine revealed a region in close proximity and medio-caudal with respect to striatum that is heavily innervated by histamine-containing axons which both anatomical analysis as well as further immunohistochemistry (e.g. labelling for CTIP2 and PKC-δ) suggests corresponds to the ovBNST (<u>Franklin, 2007</u>). Indeed, earlier studies in adult rodents did already demonstrate the existence of a significant histaminergic projection to the BNST (<u>Ben-</u> <u>Ari, 1977</u>) as well as histamine enzymatic activity in this region (<u>Cumming, 1991b</u>). More recent studies in mature transgenic mice also confirm this dense innervation of BNST by histaminergic axons (<u>Lin et al., 2023</u>). The BNST is a highly complex subcortical nucleus composed of many different subnuclei and critical in the regulation of many stress and anxiety-related behaviors (<u>Fox et al., 2010</u>; <u>Kim et al., 2013</u>; <u>Lebow & Chen,</u> <u>2016</u>; <u>Tinterri et al., 2018</u>; <u>Wang et al., 2019</u>; <u>Donner et al., 2020</u>; <u>Tsukahara & Morishita,</u> <u>2020</u>; <u>Engelhardt et al., 2021</u>) and is thought to be integral in the regulation of limbic structures and larger networks guiding autonomic behaviors (<u>O’Connell & Hofmann,</u> <u>2012</u>; <u>Albin, 2018</u>). Indeed, the BNST has been suggested to be a central hub within the so-called social decision-making network or SDMN linking the motor and reward brain networks (e.g. the basal ganglia) with those involved in guiding social behaviors (<u>Albin, 2019</u>). Our findings are in line with observations of a dense histaminergic innervation of the BNST and demonstrates that this innervation is already present and established by the start of the second postnatal week (and likely earlier) and appears to be restricted to the oval nucleus of the BNST. Activity within this nucleus is thought to promote anxiogenic behavior (<u>Crowley et al., 2016</u>; <u>Boucher et al., 2022</u>), but the vast heterogeneity of neuronal subtypes even within this single subnucleus suggest that its functional roles are likely more complex (<u>Larriva-Sahd, 2006</u>; <u>Ortiz-Juza et al., 2021</u>). Before investigating whether this nucleus could be a source of striatal histamine it was first explored to what extent striatal neurons and synapses were responsive to histamine during this developmental period.

These first experiments shows that young striatal spiny projection neurons (SPNs) and their cortical inputs can indeed both be modulated by exogenously applied histamine. Striatal SPNs consist of two different types: the dopamine-receptor 1 (D_1_) and the dopamine-receptor 2 (D_2_) expressing SPNs which correspond to the direct and indirect pathway neurons respectively (<u>Gerfen et al., 1990</u>; <u>Smith et al., 1998</u>) and selectively innervate downstream basal ganglia nuclei and differently affect motor behaviour (<u>Kravitz et al., 2010</u>). It has been shown that histamine tends to depolarize SPNs in adulthood (<u>Ellender et al., 2011</u>; <u>Rapanelli et al., 2016</u>; <u>Zhuang et al., 2018</u>; <u>Aceto et al., 2022</u>) and more recently we showed that young developing SPNs are also modulated by histamine and are either hyperpolarized or depolarized depending on their developmental age (<u>Han et al., 2020</u>). As the SPNs form these distinct functional pathways here we expanded on these earlier findings by including *posthoc* immunostaining of patched SPNs and show that in the second postnatal week both D_1_ and D_2_ SPNs in the medial striatum respond similarly to histamine superfusion. However, to our surprise histamine tended to hyperpolarize instead of depolarize the D_1_ and D_2_ SPNs as would be expected at this age (<u>Han et al., 2020</u>). Several explanations might be put forward. Firstly, recordings were restricted to dorsomedial SPNs as compared to the dorsal striatum suggesting this might be a different restricted population (<u>Shen et al., 2004</u>; <u>Han et al., 2020</u>; <u>Alegre-Cortes et al., 2021</u>). Secondly, a major difference is that the aCSF used here contained GABA receptor antagonists. As histamine negatively modulates striatal GABA release this might it other instances change the striatal GABAergic tone facilitating depolarization (<u>Ellender et al., 2011</u>; <u>Han</u> <u>et al., 2020</u>). Additionally, histamine has also been shown to directly bind and alter gating of certain extrasynaptic GABA_A_ receptors suggesting complex interactions between histamine and GABA mediated signaling (<u>Saras et al., 2008</u>; <u>Bianchi et al., 2011</u>; <u>Sente et al., 2022</u>). Modulation of other processes were consistent with previous observations and included reductions in action potential frequency, prolongation of the interspike intervals (ISI) (<u>Nisenbaum et al., 1994</u>; <u>Sittig & Davidowa, 2001</u>; <u>Baranauskas et</u> <u>al., 2003</u>; <u>Shen et al., 2004</u>; <u>Shen et al., 2005</u>; <u>Han et al., 2020</u>) and negative modulation of cortical synaptic inputs (<u>Doreulee et al., 2001</u>; <u>Ellender et al., 2011</u>; <u>Han et al., 2020</u>). Indeed, concomitant with the decrease in intrinsic excitability we find that exogenously applied histamine negatively modulates transmission at the glutamatergic inputs onto SPNs arriving from both the mPFC and visual cortex. Together these results based on exogenous application of histamine suggest it is an active neuromodulator at both the D_1_ and D_2_ SPNs during this developmental period and raises the question what the physiological sources of histamine might be in the absence of striatal innervation.

As histamine is thought to be released all along the axons coming from histaminergic neurons in the TMN this suggests that densely innervated regions can form a source of physiological histamine. Indeed, using trains of electrical stimulation of the ovBNST and using recently developed GRAB_HA_ sensors expressed in striatal SPNs we show that SPNs located hundreds of micrometers away from the ovBNST can detect significant levels of histamine (<u>Dong et al., 2023</u>). Similarly, FCV recordings in striatum made at similar distances showed that electrical stimulation of the ovBNST induced a prolonged histamine transient ranging in the tens of seconds as well as an initial fast dopamine transient. These different time frames are consistent with observations suggesting that dopamine is rapidly removed from the extracellular space (<u>Threlfell et</u> <u>al., 2012</u>; <u>Sulzer et al., 2016</u>; <u>Hamid et al., 2021</u>), whereas histamine can still be detected for prolonged periods ranging in the tens of seconds to minutes once released (<u>Biggs</u> <u>& Johnson, 1980</u>; <u>Prast, 1989</u>; <u>Mochizuki, 1991</u>; <u>Dong et al., 2023</u>) more akin to neuromodulators such as adenosine and serotonin (<u>Threlfell et al., 2004</u>; <u>Roberts et al.,</u> <u>2022</u>). This likely partly results from the kinetics of the organic cation transporter 3 (OCT3) and membrane-bound histamine N-methyltransferase (HNMT) in removing active histamine (<u>Barnes & Hough, 2002</u>; <u>Duan & Wang, 2010</u>). Indeed, the relatively long lifespan of histamine as shown by others (<u>Cumming, 1991a</u>; <u>Dong et al., 2023</u>) led us to consider that the ovBNST could act as a significant histaminergic source for the developing striatum and potentially other structures close by. To demonstrate that the levels of histamine are sufficient to modulate striatal physiology we combined optogenetic stimulation of mPFC afferents with whole-cell patch-clamp recordings of SPNs and used electrically evoked release of histamine from the ovBNST. These experiments revealed that released histamine can induce a transient reduction of glutamatergic transmission from mPFC afferents by acting at H_3_ receptors. To our knowledge this is the first direct evidence of neuromodulation and volume transmission for histamine across anatomical boundaries (<u>Rice & Cragg, 2008</u>; <u>Fuxe et</u> <u>al., 2010</u>). This is the case during early development, and it remains to be seen whether this is true also during adulthood, although the BNST is densely innervated by histaminergic afferents in adulthood also (<u>Lin et al., 2023</u>).

Overall, our results strengthen the evidence for early life modulatory effects of histamine in the striatum and provides a solution to the conundrum what the physiological sources of striatal histamine might be. To our best knowledge this is first demonstration of a dense histaminergic axonal plexus in the ovBNST capable of releasing histamine which appears to act as a paracrine neuromodulator. This provides not only a novel view of the sources and modes of action of histaminergic transmission in normal brain physiology and development, but might have implications also for the study and management of brain disorders such as Tourette’s syndrome and OCD (<u>Luyten et al., 2016</u>). Indeed, although a variety of pharmacological and non-pharmacological treatments exist (<u>Bloch, 2008</u>) in severe cases deep brain stimulation (DBS) of the BNST has been shown to be efficacious (<u>Denys et al., 2010</u>; <u>Nuttin et al.,</u> <u>2013</u>; <u>Luyten et al., 2016</u>; <u>Mosley et al., 2021</u>; <u>Naesstrom et al., 2021</u>). Although the effects of DBS are complex and exert their actions on multiple time scales, including longer term changes in prefrontal cortex connectivity to subcortical targets (<u>Figee et al., 2013</u>; <u>Baldermann et al., 2019</u>) it is tempting to speculate that release of neuromodulators such as histamine might be responsible for some of the rapid onset effects.

## Methods

### Animals

All experiments were conducted on C57Bl/6 WT mice of both sexes with *ad libitum* access to food and water. Experiments were designed to use all littermates across developmental age ranges to control for effects of litter sizes and maternal care factors that could affect degree of neuronal and circuit maturity. All mice were bred and housed in a temperature-controlled animal facility (normal 12:12 h light/dark cycles) and used in accordance with the UK Animals (Scientific Procedures) Act (1986) and European Ethics Committee (decree 2010/63/EU) and were approved by the Committee on Animal Care and Use at the University of Oxford, UK and Ethische Commissie Dierproeven at the University of Antwerp, Belgium.

### Immunohistochemistry and image analysis

The brains of P9-11 C57 mice were immersed and fixed for 3 days in cold 1-ethyl-3-[3-dimethylaminopropyl] carbodiimide HCl (EDC, Thermo Fisher Scientific, in 0.1 M PB, pH 7.4) at 4°C, followed by 4% paraformaldehyde in 0.1 M PBS for 2 days at 4°C, and kept in PBS until used. Sagittal 70 µm sections were made on a vibratome (Leica 1000S) and washed (PBS, 3x, each 10 min.). Sections underwent antigen retrieval by heating at 80°C in 10 mM sodium citrate, pH 6.0, for 30 min, after which they were washed and blocked for 1 hour at room temperature with 10% Normal Goat Serum in PBS-Tx 1%. Sections were incubated with histamine antibody (rabbit, 1:500 in PBS, Immunostar, Cat. 22939, Lot. 1532001) for 72 hours at 4°C under gentle agitation, followed by incubation overnight with biotin-XX goat anti-rabbit (1:500 in PBS, Invitrogen, Cat: B2770, LOT:1870403). To amplify the histamine signal, brain sections were incubated with Vectastain ABC kit for 4 hours at room temperature. After 2 hours of incubation with the Vectastain ABC kit, CTIP2 (chicken ovalbumin upstream promoter transcription factor-interacting protein) antibody was added (rat, 1:500, Abcam, Cat: ab18465, Lot: GR3272266-13) to facilitate delineation of striatum and incubation continued until the 4 hours were completed. Brain slices were washed as before and incubated overnight with Streptavidin-Cy3 (1:1500 on PBS, ZyMax, Invitrogen) and anti-rat Alexa Fluor 647 (1:1000 on PBS) at 4°C. When triple staining was done to also delineate the BNST, PKC-δ antibody (mouse, 1:1000 in PBS, BD Transduction Laboratories Cat: 610398) was added and incubated overnight at 4°C followed by overnight incubation with anti-rat Alexa Fluor 488. DAPI was added to sections and incubated for 10 min at room temperature (1:1000 in PBS, Sigma Cat. 11190301, Lot. 100666). Images of the double or triple stained brain section were obtained with a confocal (Olympus FV1000) or epifluorescence microscope (Olympus BX40). All images were analyzed using Fiji Image J software (<u>Schindelin et al., 2012</u>). For the images in Figure 3, CTIP2 staining was consider a suitable marker demarcating the full area of the striatum and was used to draw and divide the striatum in equally sized sections: An Upper Striatum (US) ranging from the upper limit of the striatum until the lower limit of the lateral ventricle, a Lower Striatum (LS) ranging from the limit of the lateral ventricle to the upper limit of the anterior commissure (*ac*), and a Ventral Striatum (VS) ranging from the upper limit of the *ac* to the limit of CTIP2 staining (Figure 3F). The internally built Image J Plot Profile plug-in was used to obtain a pixel intensity scan along the x-axis of the labelled brain sections. This scan compiled the pixel intensity per column (obtained as pixel intensity per μm) along the y-axis. The total Area Under the Curve of histamine staining (US + LS + VS) was obtained from the scans and normalized to obtain the relative probability of finding histamine fibers in either striatal section. For images in Figure 4, an ROI was manually drawn around PKC-δ positive cell bodies which were considered ovBNST neurons. The rostral part of the ovBNST was used to delineate the anatomical rostral location of the BNST. The area surrounding the ovBNST, was considered as BNSTvm and a second ROI was drawn to delineate this region. Both ROIs were saved and superimposed on the histamine-stained images independently from which the mean intensity of the histaminergic signal was obtained.

### Intracerebral AAV-ChR2 injections

Postnatal day (P)0-3 mice were anesthetized by isoflurane (5% induction, followed by 1% maintenance) in conjunction with brief ice-induced anesthesia (<1 min). Mice were injected using a glass pipet containing AAV1-CaMKIIa-hChR2(H134R)-eYFP (<u>Lee et</u> <u>al., 2010</u>) (Addgene) and the dye fast green (total volume ∼0.5 μl/injection) in the medial prefrontal cortex (mPFC) or the visual cortex and left to recover on a heating pad at (37° C). After this, they were placed back in the cage with the dam and other littermates.

### Brain slice preparation

Acute striatal slices were made from mice between P9–15 of age. Mice were anesthetized with isoflurane and rapidly decapitated. Coronal 350–400 µm slices were cut using a vibrating microtome (Microm HM650V). Slices were prepared in aCSF containing the following (in mm): 65 sucrose, 85 NaCl, 2.5 KCl, 1.25 NaH_2_PO_4_, 7 MgCl_2_, 0.5 CaCl_2_, 25 NaHCO_3_, and 10 glucose, pH 7.2–7.4, bubbled with carbogen gas (95% O2/5% CO2). Slices were immediately transferred to a storage chamber containing aCSF (in mM) as follows: 130 NaCl, 3.5 KCl, 1.2 NaH_2_PO_4_, 2 MgCl_2_, 2 CaCl_2_, 24 NaHCO_3_, and 10 glucose, pH 7.2–7.4, at 32°C and bubbled with carbogen gas until used for recording.

### Whole-cell patch-clamp recordings

Whole-cell patch-clamp recordings from single SPNs were made in dorsomedial striatum in acute brain slices placed in a submerged recording chamber (Slicescope Pro 1000, Scientifica) continuously perfused with aCSF (32°C and perfusion speed of 2 ml/min) with the same composition as the storage solution and bubbled with carbogen gas. As standard both gabazine (SR95531, 200 nM) and CGP52432 (1 μM) were added to aCSF to block GABA_A_ and GABA_B_ receptors respectively, and for certain experiments histamine (10 μM) or the H_3_R antagonist thioperamide (10 μM) were additionally added. Whole-cell current-clamp recordings were performed using glass pipettes, pulled from standard wall borosilicate glass capillaries (to minimize dialysis of cytosolic components 6-8 mΩ resistance pipettes were used), and containing (in mM): 110 potassium gluconate, 40 HEPES, 2 ATP-Mg, 0.3 Na-GTP and 4 NaCl (pH 7.2–7.3; osmolarity, 290-300 mosmol/L). Recordings were made using a Multiclamp 700B amplifier and filtered at 4 kHz and acquired at 10 kHz using an InstruTECH ITC-18 analog/digital board and WinWCP software (University of Strathclyde, RRID: SCR_014713) at 10 kHz.

### Optogenetic stimulation of cortical afferents

Whole-cell patch-clamp recordings were made from SPNs as standard in striatal slices of animals previously injected with AAV1-CaMKIIa-hChR2(H134R)-eYFP (<u>Lee et al.,</u> <u>2010</u>) (Addgene) in either mPFC or the visual cortex. Afferents were optically stimulated every 5 seconds for at least 5 minutes or until a stable EPSP amplitude was observed (less than 10% variation in amplitude). Photoactivation of ChR2 was achieved using widefield 2-5 ms duration 473nm blue light pulses of ∼1 mW via a TTL triggered CoolLED pE-300 system (CoolLED, Andover, UK). After a stable baseline EPSP amplitude was obtained histamine (10 μM) was added to the aCSF and superfused for at least 15 minutes after which the EPSP amplitude was assessed again.

### Recording and analysis protocols

Hyperpolarizing and depolarizing current steps were used to assess the intrinsic properties of the recorded SPNs including input resistance, spike threshold (using small incremental current steps) and membrane time-constant, as well as the properties of action potentials (amplitude, frequency, and duration). Properties were assessed immediately on break-in. Currents step ranges were -100pA to +100pA. Data were analyzed offline using custom written programs in Igor Pro (Wavemetrics, RRID: SCR_000325). The input resistance was calculated from the observed membrane potential change after hyperpolarizing the membrane potential with a set current injection. The membrane time constant was taken as the time it takes for a change in potential to reach 63% of its final value. The action potential amplitude was taken from the peak amplitude of the individual action potentials relative to the average steady-state membrane depolarization during positive current injection. Action potential duration was taken as the duration between the upward and downward stroke of the action potential at 25% of the peak amplitude. Evoked EPSPs were defined as upward deflections of more than 2 standard deviations (SD) on average synaptic responses generated after filtering and averaging original traces (0.1 Hz high-pass filter and 500 Hz low-pass filter) and used for analysis of synaptic properties. Synaptic properties include measurements of peak amplitude, duration (measured from the start of the upward/downward stroke of the event until its return to the pre-event baseline), rise time (time between 20% and 80% of the peak amplitude) and decay time (measured as the time from peak amplitude until the event returned to 50% of peak amplitude). The effect of electrical stimulation of ovBNST on optogenetically evoked EPSPs was studied by optically eliciting EPSPs every 5 s. for at least 5 min. to obtain a stable baseline response. Following this the ovBNST was electrically stimulated at 5 Hz for 3 min. after which EPSPs were optically elicited for a further 10 min. EPSPs were normalized to the average EPSP amplitude of the last minute of baseline. The barplots in Figure 6D depict the baseline EPSP amplitude and the average EPSP amplitude 4-5 minutes after electrical stimulation of the ovBNST (average of 12 optically evoked EPSPs) in normal aCSF and aCSF containing thioperamide.

### Fast scan cyclic voltammetry (FSCV) experiments

Acute brain slices were generated and stored as standard (see above) until used. Carbon fiber microelectrodes (CFMs) were made in-house as previously described (<u>Abdalla et al., 2020</u>). Briefly, a single carbon fiber (7 μm, Goodfellow Corporation, Coraopolis, PA, USA) was aspirated into a glass capillary (0.6 mm OD, 0.4 mm ID, 10 cm long; A-M Systems, Sequim, WA, USA) and pulled under gravity and heat with a vertical pipette puller (Narishige Group, Tokyo, Japan) to form a carbon-glass seal. The exposed carbon fiber was cut to ∼150 µm under a light microscope. An electrical connection was made with the fiber by inserting a stainless-steel wire (Kauffman Engineering, Cornelius, OR, USA) coated with silver epoxy into the capillary. To maximize sensitivity to dopamine and histamine, the carbon surface was modified two ways: first, a thin layer of Nafion^TM^ (Ion Power, New Castle, DE, USA) was electrodeposited at 1 V for 30 s, and cured at 70⁰C for 10 minutes (<u>Hashemi et al., 2009</u>), then glutamic acid was electropolymerized using a polymerization waveform (-1.0 to 1.2 to -1.0 V at 400 V/s applied at 60 Hz) for 10 min (<u>Holmes et al., 2021</u>).

All FSCV calibrations were performed in recording aCSF bubbled with carbogen gas for at least 30 min prior to measurements. CFMs were calibrated in a flow cell with known concentrations of dopamine and histamine. Dopamine and histamine were dissolved in recording aCSF bubbled with carbogen for at least 30 min prior to experimentation. Serial dilutions were performed to create calibration solutions of dopamine (1, 0.5, 0.25, 0.1 and 0.05 μM) and histamine (5, 1, 0.5, 0.25, 0.1 µM). Flow injection analysis (FIA) was performed in a custom-built flow cell (<u>Hexter et al.,</u> <u>2023</u>), with a syringe infusion pump (Harvard Apparatus, model 70-4500, Cambridge, UK) controlling the flow at 1.7 mL/min. A pseudo-Ag/AgCl reference electrode was placed below the CFM and the hole was plugged to seal the system. The CFM was mounted and lowered into the flow stream until the exposed fiber was fully submerged. The histamine-optimized waveform (-0.5 to -0.7 to 1.1 to -0.5 V at 600 V/s) was applied at 60 Hz for 10 min and then at 10 Hz for 10 min prior to data collection. For each calibration point, the analyte was introduced to the flow stream for 10 s via a six-port HPLC loop injector (Cheminert valve, VIVI, Houston, TX, USA) resulting in a rectangular plug. Each point was repeated three times per electrode.

Brain slices containing the BNST and striatum were placed into a bath of recording aCSF bubbled with carbogen gas at 30 ⁰C flowing at 2 mL/min with a peristaltic pump. The slice was secured with a harp with nylon threads. Recordings with CFMs were made in striatal slices in a submerged recording chamber (as used for whole-cell patch-clamp recordings) continuously perfused with aCSF (32°C and perfusion speed of 2 ml/min) with the same composition as the storage solution and bubbled with carbogen gas. The CFM was then placed in the striatum and allowed to equilibrate with the histamine waveform applied at 60 Hz for 10 minutes then 10 Hz for 10 minutes. Electrical stimulation was applied (20 Hz, 30 pulses at 120-200 µA) to the ovBNST using a twisted tungsten stimulating electrode, placed with an identical quadrupole micromanipulator.

FSCV data was acquired using WCCV 4.0 software (Knowmad Technologies, LLC, Tuscon, AZ, USA), a USB-6341DAC/ADC (National Instruments, TX, USA) device, a Dagan potentiostat (Dagan Corporation, Minneapolis, MN, USA) and a Pine Research head stage (Pine Research Instrumentation, Durham, NC, USA). Histamine and dopamine were co-monitored using a histamine-optimized waveform (-0.5 to -0.7 to 1.1 to -0.5 V at 600 V/s) at 10 Hz (<u>Samaranayake et al., 2015</u>) during and immediately after electrical stimulation. All FSCV measurements are reported against pseudo-Ag/AgCl reference electrode made prior to experimentation by applying 5 V across two silver wires (A-M Systems, Sequim, WA, USA) in 1 M HCl for ∼30 s to deposit silver chloride onto the surface.

All data collected at the CFM were filtered in WCCV using a 4^th^ order Butterworth low-pass filter with a cut-off frequency of 3 kHz and smoothed twice. To get rid of any electrochemical drift, all data were filtered in MatLab (2^nd^ order Butterworth bandpass filter with cutoff frequencies of 0.01 and 1.5 Hz). Data are reported as the average ± standard error of the mean (SEM).

### GRAB_HA_ fluorescent reporter imaging

*In utero* electroporation (IUE) of the lateral ganglionic eminence (LGE) was performed using standard procedures (<u>van Heusden et al., 2021</u>) to express the histamine-sensitive fluorescent reporter GRAB_HA_ in striatal SPNs. In short, pregnant females were anaesthetized using isoflurane and their uterine horns were exposed by midline laparotomy. HA1h GRAB_HA_ plasmid DNA (∼3.0 μg/μl), in which expression of the fluorescent histamine GRAB sensor is under the control of the hSyn promoter (<u>Dong</u> <u>et al., 2023</u>), combined with 0.03% of the dye fast green was injected intraventricularly using pulled micropipettes through the uterine wall and amniotic sac. Total volume injected per pup was ∼1 μl. Typically, around 80% of the pups underwent electroporation. Afterwards the uterine horns were placed back inside the abdomen, the cavity was filled with warm physiological saline and the abdominal muscle and skin incisions were closed with vicryl and prolene sutures, respectively. Dams were placed back in a clean cage and monitored closely until the birth of the pups. Acute coronal striatal slices were made from electroporated mice between P9–15 as outlined above. Slices were moved to a submerged recording chamber (Slicescope Pro 1000, Scientifica) continuously superfused with aCSF (32°C and perfusion speed of 2 ml/min) and striatal SPNs expressing GRAB_HA_ in proximity to the ovBNST were targeted based on the endogenous eGFP expression. Electrical stimulation to the ovBNST was performed using a twisted tungsten stimulating electrode (30 sec., at 5 Hz with 100 μA). Before and during stimulation the fluorescence was measured from neurites of GRAB_HA_-expressing striatal SPNs using brief 1-2 sec. long 473 nm wide-field illumination CoolLED pE-300 system (CoolLED, Andover, UK) combined with image acquisition (at 10 Hz) using a Hamamatsu ORCA camera and HCImageLive software. Video files were analyzed with the Sequence Intensity Analysis function within the HCImageLive software by placing multiple circular regions of interest (ROIs) on neurites of GRAB_HA_ expressing SPNs. The fluorescence intensity from these ROIs were measured and were used to derive a background-subtracted fluorescence intensity (*Ft*) by subtracting average background fluorescence intensity from equal-sized ROIs where there was no GRAB_HA_ expression (i.e., in between neurites). Data are expressed as a change in fluorescence (Δ*F/F_0_*) and were derived by calculating [(*F_t_* – *F_0_*)/*F_0_*], where *F_0_* is the average *F_t_* of the first 10-20 acquired images (initial 1-2 sec.). The final 1-2 sec. into the 5 Hz stimulation train were used to acquire images (10-20 images, acquisition rate 10 Hz) reflecting extracellular histamine levels in response to the stimulation and the maximum Δ*F/F_0_* is plotted. Data analysis of bath application of exogenous histamine (10 uM) was similar and (*ΔF*/*F_0_*) obtained 5 minutes into superfusion of histamine.

### BNST stimulation and modulation of cortical transmission

Sagittal brain slices (80-100 μm, Microm HM650V) were obtained from P9-12 C57Bl/6 mice injected in the medial prefrontal cortex (mPFC) at P0-3 with AAV1-ChR2-eYFP as detailed above. Thin brain sections allowed for an increased number containing the BNST. Brain slices were checked for the anatomical landmarks including the anterior commissure (*ac*) and anterior arm of the anterior commissure (*aca*; Figure 5) indicating the location of ovBNST. Once brain slices were selected, they were transferred to a Petri dish containing cutting aCSF, placed under a microscope and very gently a cut was made from the ventricle to the lower border of the ventral striatum (Figure 5), passing between the *ac* and the *aca,* to sever all direct axonal connections between BNST and striatum. The cut brain slices were used for experiments and were superfused with aCSF containing SR95531 (200 nM) and CGP52432 (1 μM) at a low circulation speed (5 rpm). An electrical stimulation electrode was placed above the *ac* in the ovBNST containing histamine fibers and during placement the tissue was moved slightly closer towards striatum if separated by the cut. Whole-cell patch-clamp recordings of SPNs were obtained as outlined above. EPSPs were obtained by optogenetic activation of ChR2 expressing axons coming from mPFC every 5 seconds for a total period of at least 5 minutes to form a baseline (CoolLED, Andover, UK). After this, the ovBNST was electrically stimulated for 3 min, at 5 Hz and ∼100 μA strength, to induce histamine release from the ovBNST, followed by 10 minutes recording optically evoked EPSPs every 5 seconds. When the involvement of histamine H_3_ receptor was tested, the H_3_R antagonist thioperamide (10 μM) was added to aCSF prior to recordings.

### Statistics

All data are presented as means ± SEM. Statistical tests were all two-tailed and performed using SPSS 17.0 (IBM SPSS statistics, RRID:SCR_002865) or GraphPad Prism version 9.0 (GraphPad software, RRID:SCR_002798). Continuous data were assessed for normality and appropriate parametric (ANOVA, paired *t*-test and unpaired *t*-test) or non-parametric (Wilcoxon) statistical tests were applied (* p<0.05, ** p<0.01, *** p<0.001).

## Acknowledgements

We would like to thank all members of the Ellender lab for advice and comments and Yulong Li for the kind sharing of GRAB_HA_ plasmids. Some graphics were generated using BioRender.com.

## Data sharing plans

All data is freely available – please contact corresponding author for information.

